# Label-free phenotyping of human microvessel networks

**DOI:** 10.1101/2024.02.20.581133

**Authors:** Luca Rappez, Akinola Akinbote, Marta Cherubini, Virginie Uhlmann, Kristina Haase

## Abstract

Understanding the spatial heterogeneity in blood vessel formation and development is crucial for various biomedical applications. Traditional methods for *in-vitro* microvessel segmentation rely on fluorescent labeling, which either interferes with the sample homeostasis, limits the study to a restricted set of precursor cells, or requires sample fixation, thus preventing live measurements. Moreover, these methods often focus on small, cropped images, neglecting global spatial heterogeneity of microvasculature, leading to biased data interpretation. To overcome these limitations, we present VascuMap, a deep-learning-based tool for label-free vessel segmentation and spatial analysis. VascuMap enables a comprehensive examination of entire vessel networks, capturing both morphological and topological features across the full vascular bed. Our method achieves high segmentation accuracy, comparable to the state-of-the-art fluorescence-based models. VascuMap’s capabilities extend to characterizing vasculature generated from label-free patient-derived samples, a vital step towards personalized medicine. Its compatibility with widefield label-free microscopy also accelerates sample acquisition, making it ideal for high-throughput systems crucial for drug toxicity and safety screens.

## Introduction

Historically, vascular research predominantly focused on larger blood vessels, mainly due to ease of imaging and physical accessibility. In contrast, microvessels received less attention, despite their critical functions in regulating nutrient and gas exchange^1^, facilitating immune cell activity^2^, and their significant implications in various diseases^3^. The recent introduction of microfluidic models - referred to as microvessel-on-chip - has allowed for comprehensive studies into human-derived microvessels for the first time^4^. These models have shed light on processes such as angiogenesis^5^, vasculogenesis, tumor cell extravasation and immune cell dynamics^6,7^, and vascular barrier function under varied environmental contexts. Exploiting the ability of endothelial cells to self-organize, microphysiological systems have revealed the complex, heterogeneous nature of microvessels^4^.

Vascular beds across all tissues inherently exhibit a complex spatial heterogeneity, a phenomenon also evident in microvessel-on-chip experiments^8^. This heterogeneity stems from various factors, including intricacies in cell communication during the formation of blood vessels, the microvasculature’s responsiveness to mechanical forces, local disparities in the concentration of the extracellular matrix (ECM), differential ECM breakdown, and uneven flow within and around the vessels^9,10^. Such dynamics play a pivotal role in the remodeling of these networks. Consequently, we observe significant spatial variability within and across microvascular samples *in vitro*, mirroring the patterns seen in living organisms.

To effectively investigate microvascular physiology, it is therefore essential to characterize the network in its entirety. This requires accurate segmentation of microscopy data to extract quantifiable morphological, topological and spatial features of vessel networks. Presently, segmentation of *in-vitro* human microvasculature is exclusively reliant on fluorescent labeling techniques such as dyes, antibody-based labeling, or genetic integration of fluorescent reporters in a subset of precursor cells^11–16^. Although such methods yield a varied set of morphometrics associated with various vascular drug responses^17,18^, they either limit the analysis to fixed and permeabilized samples or disrupt sample homeostasis, ultimately impeding the live examination of unaltered samples over time. Furthermore, current techniques often omit spatial heterogeneity, as they focus on segmented cropped areas rather than the entire vascular network^19^. Doing so ultimately leads to a skewed understanding of the complex heterogeneity across and within individual vascular networks, as interpretations are biased by the limited fields of view selected from the sample cohort.

To jointly address these limitations, we introduce VascuMap, a deep-learning-driven approach for label-free vessel segmentation and spatial characterization. VascuMap allows for a comprehensive assessment of the entire vessel network, accounting for spatial heterogeneity by inspecting the combined morphological and topological features of the network. Our approach delivers high segmentation accuracy from the brightfield modality and is comparable to models trained on fluorescence-segmented vessels, thereby showcasing its efficacy. By employing graph-based topological metrics as well as morphological features to categorize vessels cultured under different conditions, we demonstrate the ability of VascuMap to capture nuanced physiological variations.

We expect VascuMap to facilitate the characterisation of label-free patient-derived microvessels, a critical step in the development of preclinical tools for personalized medicine or safety testing of pharmaceuticals. Its compatibility with widefield microscopy in brightfield significantly speeds up sample acquisition, making it suitable for high-throughput systems, which are instrumental for drug development and testing. Additionally, the brightfield modality allows for non-invasive and long-term time-resolved analysis, thus presenting exciting venues for investigating human microvasculature development *in vitro*.

## Results

Herein, we introduce VascuMap, an analytical approach for a comprehensive examination of *in vitro* vascular systems in a spatial context (**Figure 1A**). This method is applied on microscopy images of *in vitro* vessels that are generated in a microfluidic chamber using endothelial cells grown together with stromal cells within a fibrin hydrogel. These self-assemble to form a perfusable microvessel network over several days. Following the formation of a microvascular bed, a three-dimensional brightfield microscopy dataset is acquired across the whole chip encompassing the vascularized microtissue. Due to the planar nature of microvessels and the misalignment with the microscope stage focal plane, microvessels appear at different heights, preventing the observation of the complete vascular network in a single 2D microscope acquisition. In addition, as optical properties of the ECM are complex and heterogeneous, existing common Z projections are not suitable^20^. Consequently a custom virtual refocusing technique was developed to measure the tilted vessel plane within the z-stack (**Supp. Fig. 1**). To achieve this, focus estimation points are set up in a grid layout on the horizontal plane of the image volume. At each grid point, 200×200 pixel patches are defined for each Z plane. The focus level of each patch is determined using the tenengrad variance, an established method^21^ which measures the variance of the image gradient amplitude, resulting in a focal score. The Z index with the highest tenengrad variance indicates the most focused plane for that area. After analyzing all focal planes across the grid, a 3D plane is fitted based on these values to approximate the overall tilted focal plane. Finally, the original Z-stack is virtually refocused along this tilted plane, producing a 2D image that displays the vessels, in focus, across the entire sample area.

**Figure 1:**
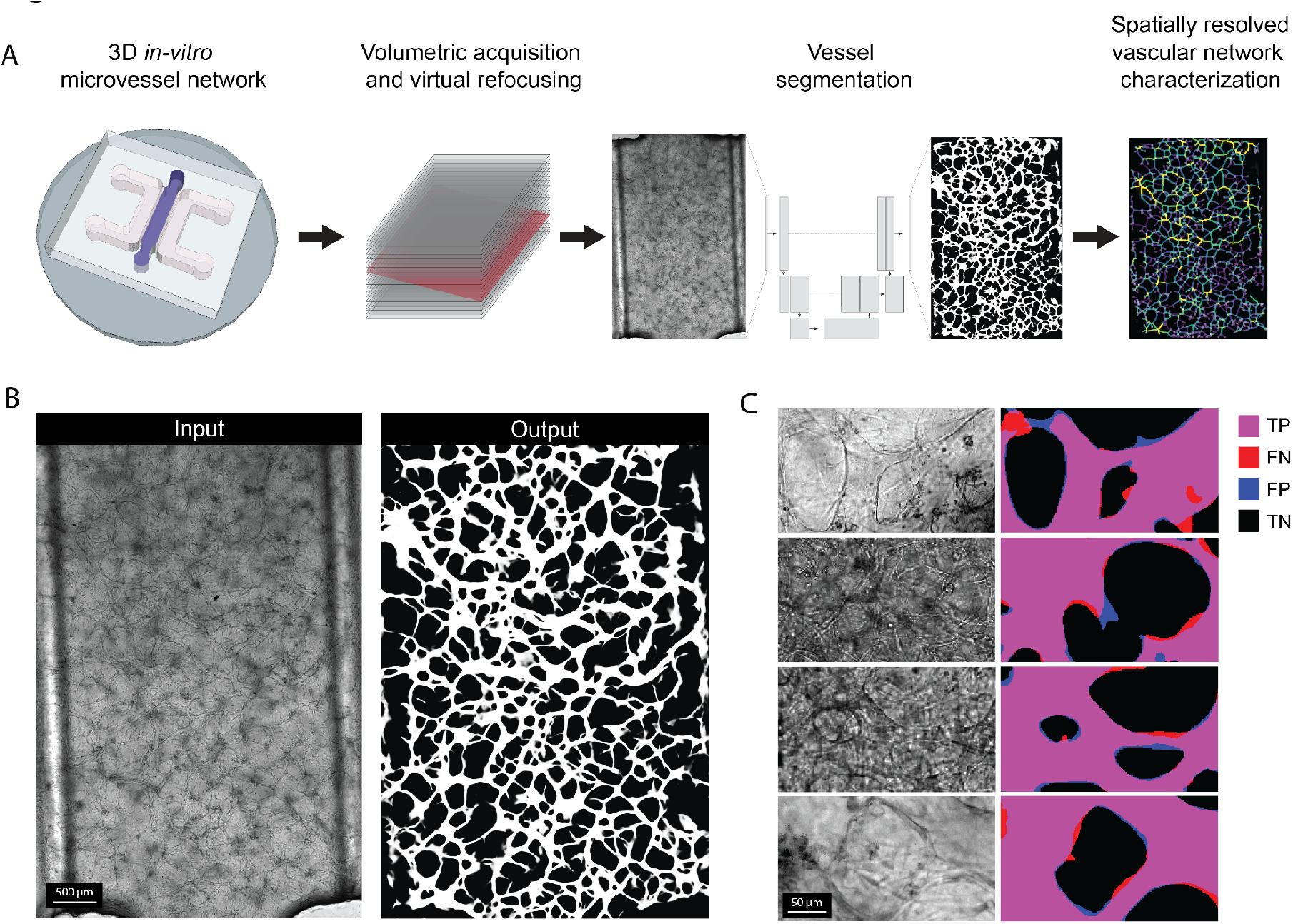
Label-free segmentation of complete vascular networks. **A**. Microvessels are cultivated within a microfluidic chamber, and a 3D Z-stack is collected in the brightfield channel. Subsequently, a 2D in-focus representation of the complete vascular network is generated through virtual refocusing. From this planar representation, vessels are segmented using a convolutional neural network. In order to provide a comprehensive analysis of the network morphology and topology, a graph is generated from the vessel mask in which the nodes represent branching points and the edges correspond to vessel pathways. **B**. Illustration of the challenges associated with segmentation of vessels in brightfield such as crowding of confounding signals and high contrast heterogeneity. **C**. Comparison of vessel segmentation on confocal (top two rows) and widefield data (bottom two rows), taking brightfield images as input (first column) and generating mask as output (second column - TP, true positives; FN, false negatives; FP, false positives; TN, true negatives).

From this refocused 2D image, the vessels are segmented using a U-Net convolutional neural network (CNN)^22^. Segmenting vessels from brightfield poses significant challenges, due in part to the confounding signal originating from other co-cultured cell types (stromal cells herein) and the complex ECM, whose protein composition is heavily remodeled during the process of vessel formation, as we have seen previously by mass spectrometry analysis^23,24^. As a result, heavy contrast variations due to differences in light absorbance across the ECM result in obscuring small vessels and some vessel boundaries (**Figure 1B-C**). Despite these challenges, we found that segmentation from fluorescence, the state of the art approach, improves by a mere 2% in accuracy over the brightfield modality, thus supporting brightfield as an appropriate modality for vessel segmentation (**Supp. Fig 2**). Further, we show that microvessels can be isolated from complex backgrounds as well as accurately interpolate these obscured small vessels (**Supp. Fig. 3A-B**). To further allow the detection of such vessels, we applied hysteresis thresholding to the pixel probability map output by the final layer of our CNN. This enables us to categorize lower-probability pixels as vessels if they are connected to high-probability pixels. This method resulted in the successful capture of small vessels and overall improved segmentation accuracies compared to a single-level thresholding method (**Supp. Fig. 3C**).

Next, we aimed to fully characterize the vascular network throughout the entire chip. We created a graph from the vessel mask, positioning nodes at vessel junctions and using edges to represent the vessels themselves (**Figure 2A**). This graph enables the calculation of diverse metrics, such as vessel length, area, tortuosity, diameter, fractal dimension, branching density, and the number of branching neighbors (**Table 1**). Together, these metrics provide a comprehensive morphological and topological insight into the vascular network. To underscore the biological significance of these metrics in highlighting physiologically pertinent traits, we performed a simple experiment to culture HUVEC microvessels (as our lab has done previously) in 2 distinct growth medias: a complete culture medium (complete condition), enriched with growth factors known to yield larger vessels and a basal minimal medium with only amino acids, glucose and ions (basal media condition), lacking additional growth factors and reduced FBS content of 2% (**Figure 2B**). Analysis using VascuMap revealed pronounced variations in the collected metrics based on these conditions (**Figure 2C**). Specifically, the complete media condition resulted in larger and longer vessels but also vascular networks with more intricate interconnections, leading to an elevated fractal dimension, which serves as an indicator of the network’s complexity. In opposition, the vessels from the basal condition were thinner and shorter, leading to a reduced overall coverage of the culture area. Consequently, the extravascular spaces between the vessels were not only larger but with a higher heterogeneity in their size and distribution, as reported by a high lacunarity value, which provides insights into the texture and spatial organization of the vascular network. This would potentially imply that the vessels from the basal condition are either still in an earlier developmental stage or in a regressed form compared to the vessels from the complete condition, as suggested by an elevated number of vessel fragments in the basal condition.

**Table 1:** Description of VacuMap’s metrics.

| Metric Name | Metric Category | Metric Type | Description |
| --- | --- | --- | --- |
| Explant area | Morphological | Global | Total area of the vessel network culture, represented by the area of a convex hull around the vascular network's mask. |
| Vessel density | Morphological | Global | Number of vessels divided by the explant area. |
| Total vessel length | Morphological | Global | Cumulative length of all the vessels in the network. |
| Total vessel area | Morphological | Global | The area occupied by the entire vessel network. |
| Total number of vessels | Topological | Global | Total count of vessels in the network. |
| Total number of sprouts | Topological | Global | Number of vessels with only one branching point. |
| Total number of branching | Topological | Global | Total number of branching points. |
| Fractal dimension | Topological | Global | Describes the complexity of the vascular network by quantifying how detail changes with scale using the box-counting method. It is estimated by performing a linear regression on the log-log plot of the number of squares - or boxes - required to cover the vessel skeleton at various box sizes. The slope of the fitted line corresponds to the fractal dimension. |
| Lacunarity | Topological | Global | Describes the texture of the vascular network, indicating gaps or holes. Similar to the fractal dimension, the lacunarity relies on the box-counting method at various box scales. It is defined as the variance-to-mean-squared ratio of non-zero elements in the box-counting histogram, averaged over scales. |
| Mean vessel diameter | Morphological | Local Vessel | The diameter of a vessel is defined as the average pixel value from its path sourced from the diameter image. The latter is obtained by multiplying the distance transform of the vessel mask by two to obtain the diameters from the radii values. |
| Mean vessel area | Morphological | Local Vessel | The sum of pixel values along a vessel path sourced from the diameter image. |
| Mean vessel orientation | Morphological | Local Vessel | The angle of individual vessels relative to the xy-coordinate plane. |
| Mean vessel aspect ratio | Morphological | Local Vessel | The ratio between a vessel's major and minor axis lengths. |
| Mean vessel tortuosity | Morphological | Local Vessel | The deviation of a vessel from a straight path represented by the ratio of the vessel length to its shortest path. |
| Mean vessel length | Morphological | Local Vessel | The sum of the Euclidean distances between adjacent coordinates along the vessel path. |
| Mean vessel shortest path | Morphological | Local Vessel | The Euclidean distance between the two extremities of a vessel. |
| Mean number of branching neighbors | Topological | Local Branching | The average number of neighboring branching points within a given radius. |
| Mean branching distance to nearest neighbor | Topological | Local Branching | The distance from a branching point to its closest neighboring branching point. |
| Mean number of vessels per branching | Topological | Local Branching | The average number of vessels connected to a branching point. |
| Mean vessel branching angle | Morphological | Local Branching | The minimum angle at which vessels branch off. |

**Figure 2:**
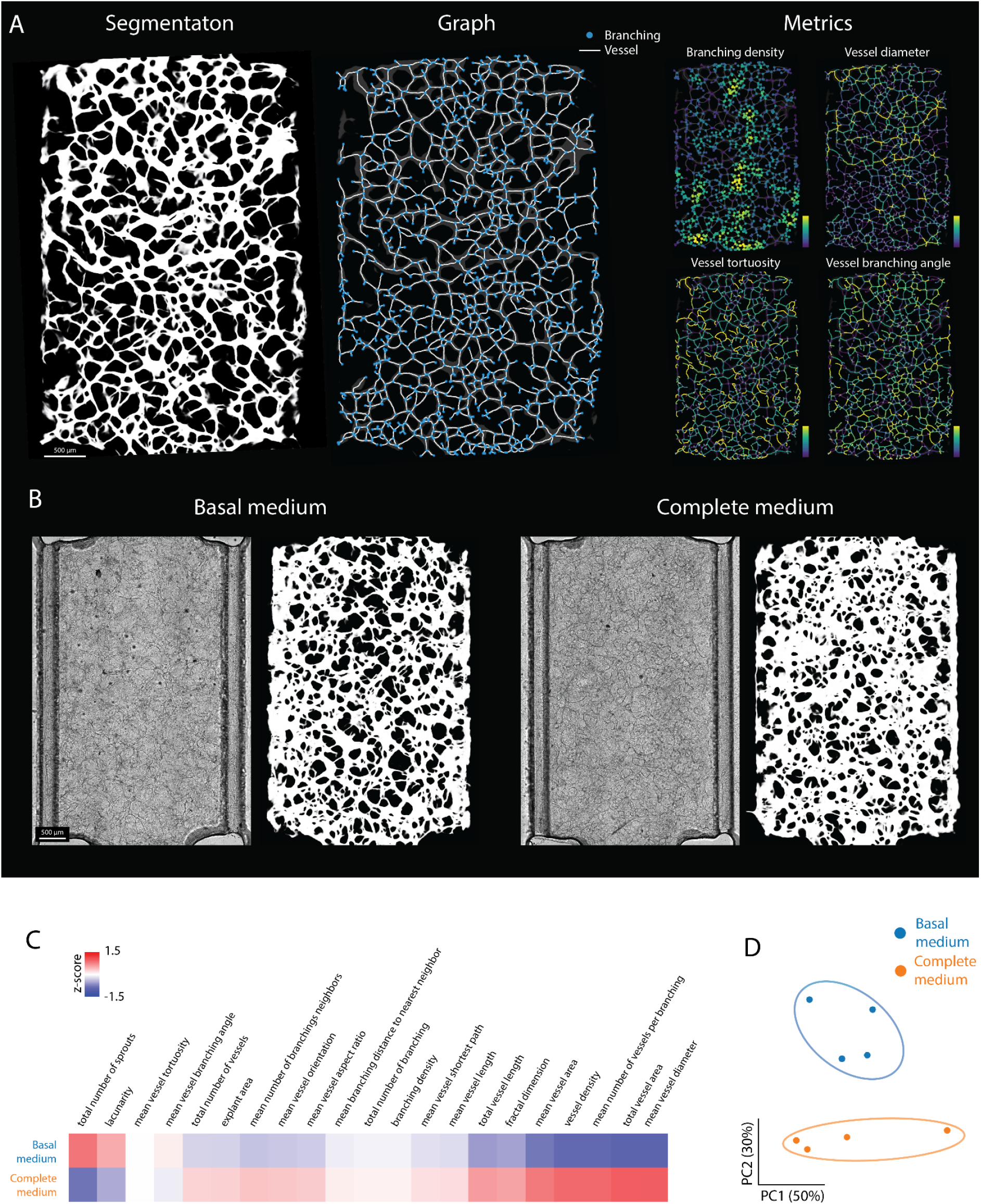
VascuMap enables spatially resolved analysis of vessel networks. **A**. Depiction of the methodology for assessing both morphological and topological features of the vascular network by constructing a graph, where nodes represent junction points and edges correspond to vessel pathways. **B**. Illustration of the microvascular networks of HUVEC endothelial cells grown in basal minimal and complete media. **C**. Heatmap showcasing variations of the metrics across the vessels grown in the two culture conditions. The color-coded values are standardized feature-wise (z-scores) and are clipped between -1.5 and 1.5 for visualization. **D**. Principal Component Analysis (PCA) constructed from the topological and morphological features, where each point symbolizes a unique vascular network sample, color-coded according to its culture condition.

Further, to demonstrate that the gathered metrics collectively characterize and distinguish the conditions, we conducted a PCA analysis (**Figure 2D**). Technical replicates of vessels cultured in Basal and complete media distinctly cluster by growth condition, showing no overlap. The lack of overlap between these two conditions reinforces the notion that the comprehensive set of features collected by VascuMap is capable of capturing biologically relevant differences in a nuanced manner.

## Discussion

In this paper, we introduce VascuMap, a comprehensive tool for segmenting and analyzing entire vascular networks spatially. Utilizing deep learning algorithms, we achieve non-fluorescent segmentation accuracies comparable to those obtained through fluorescence-based methods. This observation proves that label-free segmentation is both feasible and reliable while avoiding likely physiological alterations caused by labeling techniques, such as changes in cell viability, growth interference, and disruption of signaling pathways. By mapping a graph onto the segmented vessel mask, we jointly group morphological and topological metrics to provide an exhaustive characterization of vascular networks. We validate the utility of VascuMap and its metrics by examining the physiological changes induced by the treatment of microvessels to two different culture conditions. It is well-known that growth factors including VEGF and FGF are key regulators of sprouting angiogenesis and we have also shown their role in shaping the vasculogenesis-like processes in our microvessels on-chip. For this reason, we compared microvessels grown in a complete medium, supplemented with these and other growth factors, which promotes the formation of large, interconnected vessels and a basal minimal growth medium, serving as a factor-free reference for baseline vascular growth. The collected metrics, as well as subsequent multi-dimensionality reduction, successfully distinguish these conditions in an unsupervised manner, underscoring VacuMap’s ability to capture physiologically relevant features.

We anticipate that future research will further enhance the spatial analysis of human microvasculature. This could involve adopting methodologies from diverse fields like ecology and sociology to enable a more in-depth and quantitative study of metric co-occurrences. Another promising avenue lies in the application of specialized 3D Convolutional Neural Networks (CNNs) to label-free 3D segmentation as well as graph neural networks to offer more nuanced insights into variations in network morphology and topology that are potentially missed by the current hand-crafted metrics. In conclusion, we believe we have opened up an exciting new direction in the study of vascular generation by emphasizing a label-free approach, and we anticipate its extension to the analysis of vascular images from other species and modalities, such as through human micro-CT or imaging studies in chick and zebrafish models.

## Methods

### Cell culture

Human umbilical vein endothelial cells (HUVEC) were purchased from Lonza and cultured in endothelial media (VascuLife, Lifeline cell systems) on T75 or T150 flasks coated with 50 μg/ml rat tail collagen I (Roche). HUVEC were transduced to stably express cytoplasmic RFP (LentiBrite RFP Control Lentiviral Biosensor, Millipore Sigma Aldrich), and were used between passages 6 and 9. Normal human lung fibroblasts from Lonza (unlabelled) were cultured in the same manner as HUVEC, with the exception of the media (FibroLife, Lifeline cell systems). All cells were cultured under normal conditions at 37 °C and 5% CO2 in a standard incubator. Dissociations were carried out using TrypLE Express (Gibco), and media was completely refreshed every other day.

### Device fabrication

Devices were fabricated as previously described^25^. Briefly, PDMS (SYLGARDTM 184 Silicone Elastomer Kit, Dow) was mixed at a 10:1 elastomer to cross-linker ratio according to the manufacturer’s protocols, degassed, and poured onto a mold (arrays of chips). Following further degassing, PDMS was placed in a 60° C oven overnight and single devices were cut, punched using 1- and 2-mm diameter biopsy punches for the gel and media ports, respectively, and air-plasma bonded (Harrick systems) to clean glass slides. Assembled devices were then incubated at 60° C overnight to restore the native hydrophobic state. All devices were sterilized under ultraviolet light for at least 30 min prior to cell seeding.

### Device seeding and microvessels formation

Fibrinogen derived from bovine plasma (Sigma) was reconstituted in phosphate-buffered saline (PBS) to a working concentration of 6 mg/ml before use. Thrombin (Sigma) stock solution (100 U/ml in 0.1% w/v bovine serum albumin solution) was diluted to a 4 U/ml working solution in cold VascuLife medium. Endothelial cells and stromal cells were cultured until near-confluence before detachment and resuspended in thrombin, in separate microfuge tubes, to concentrations of 24 million endothelial cells/ml and 4.8 million stromal cells/ml. Cell suspensions were mixed 1:1 by volume and then combined with fibrinogen solution to produce a final concentration of 6 million endothelial cells/ml and 1.2 million fibroblasts/ml, in a 5:1 ratio within fibrin (3 mg/ml). The cell-gel mixture was injected into the device’s central channel and allowed to polymerize for ∼10 min at 37 °C in the incubator. Culture media was added to each media channel and devices were cultured under static conditions. The media was replaced daily until day 7. After which, the devices were rinsed with fresh PBS and incubated with paraformaldehyde 4% (PFA, Sigma) at 4°C overnight. After washing the PFA with fresh PBS, devices were stored at 4°C until the microscopy acquisition. Cultured vessels under different conditions. Basal minimal growth media (Basal VascuLife media without growth factors and reduced FBS content to 2%) was used as a control.

### Imaging

Whole devices (approximately 4×5×3mm, WxLxH) z-stacks (z-step of 5um) were either acquired in brightfield and in the RFP channel (575 nm) at 10x magnification (HC PL APO CS2 10x/0.40 DRY) on a Thunder Imager (Leica) microscope or in the transmitted light and in the RFP channel (580 nm) on a Stellaris 8 microscope using the LAS X software in both cases (Leica).

### Virtual refocusing

Due to their planar structure and alignment discrepancies with the microscope stage’s focal plane, microvessels often appear at varying heights in each plane, posing a challenge for complete visualization of the microvessel network in a single 2D acquisition (**Supp. Fig. 1**). Consequently, a 3D brightfield microscopy dataset of the entire vascularized microtissue within the chip was captured. While existing methods address the virtual refocusing of Z stacks from samples with non-planar complex topologies, these methods are either not adapted or overly complex for our near-planar microvessels^26–28^. Consequently, a tailored virtual refocusing technique was developed to accurately determine the angled plane of the vessels within the z-stack. First, focus estimation points are established in a grid layout on the horizontal plane of the image volume. For each Z plane at these grid points, 200×200 pixel patches were defined. Under the assumption that an image in focus has a greater sharpness than its out-focus counterparts, each patch’s sharpness was measured using the tenengrad variance method, which computes the variance in horizontal and vertical gradient amplitude of the image. The Z plane with the highest tenengrad variance score was identified as the most in focus for that specific patch. After analyzing all possible focal planes within the grid, a three-dimensional plane was constructed based on these scores to approximate the vessels’ overall tilted focal plane using a least-squares minimization approach. Subsequently, the original Z stack was virtually refocused, sampling pixels along this tilted plane to produce a single in-focus image of the entire microvessel network.

### Training set generation

Microscopy images of microvessels, combined from brightfield and fluorescent channels and adjusted for contrast, were uploaded into the ‘sketchbook’ iPad app. Ground truth labels were manually drawn using an Apple Pencil on an iPad 11 Pro. A layer was added over the composite image where masks were depicted in solid green. These drawings were saved as RGB .tiff files, which were then binarized using a custom Python script. First, images were turned single-channel by summing projections channel-wise, then binarized to distinguish vessels from the background. Using the Napari platform for Python, the superimposed masks on input images were inspected to identify potential labeling errors. The brightfield and fluorescence microscopy images, along with the masks, were split into 256×256 pixel tiles without overlap. These tiles were stored in distinct folders—brightfield, fluorescence, and mask—with matching filenames. Of the 1,317 manually labeled tiles, 120 were blanks (images without vessels). A total of 385 images were acquired using Leica Thunder Imager and 812 using transmitted light from the Leica Stellaris 8 microscope.

### Network training

Image patches were divided into a 70% training set and a 30% validation set. Both training and inference utilized PyTorch for deep learning model management and the Catalyst library for higher-level operations. We built multiple Unet models with different encoder architectures using the ‘segmentation-models-torch’ Python module. After evaluating various encoder architectures, the one with top validation accuracy was chosen for final model training. In our tests, the ‘mit_b5’ encoder, a mixed vision transformer, achieved the best segmentation accuracy and was selected for mask generation. This encoder, pre-trained on ImageNet, had its input layer’s three-channel weights summed to handle single-layer image input. Images underwent augmentations such as vertical/horizontal flips, translations, rotations, and scaling with mirror padding. Each model underwent 200 epochs of training with a batch size of 16, using the RAdam optimizer, an initial learning rate of 1e-3, and a step scheduler that halved the rate mid-training. The loss function is a linear combination of the binary cross-entropy and the dice coefficient at equal weights. The intersection over union (IoU) metric of the validation set was monitored at each epoch and the model achieving the best score during training was kept for inference. All models reached convergence at around 100-150 epochs as the validation loss curves reached a plateau.

### Network inference

Model inference was performed on a tiled 2D microscopy dataset covering the whole sample. As the dimensions of the tiled images differ from patch dimensions, the overlapping patch inference functionality of the segmentation-models-torch module was employed. The input image is divided into overlapping patches, which are individually processed and reassembled, with overlapping regions averaged to form the final vessel probability map. To improve results consistency, we employed the ‘flip_transform’ test-time-augmentation method from the ‘ttach’ module, which averages the predictions over multiple augmentations. To obtain the binary mask, the vessel probability maps were binarized using a hysteresis thresholding approach, with thresholds set at 0.15 (lower) and 0.5 (upper) (**Supp. Fig. 3**).

### Graph analysis

Starting from the binary image of the vascular network, a skeletonized representation was generated to reduce the vessel structures to single-pixel widths while preserving their topology. The skeletonized image served as the basis for constructing a graph using the ‘sknw’ library, which identified nodes at vessel junctions and edges representing the vessels themselves. This graph was then converted into a NetworkX graph object for further analysis. Each node and edge was annotated with spatial coordinates. A detailed list of the metrics as well as their implementation is provided in **Table 1**.

## Author contributions

L.R. and K.H. conceived the method. L.R. developed the method. L.R, K.H, A.A. and M.C. performed the experiments and collected the data.. L.R., A.A., K.H and V.U. interpreted data. L.R., K.H, A.A, V.U. and M.C wrote the paper. L.R. and K.H. supervised and coordinated the work.

## Supplementary Figures

**Supplementary Figure 1:**
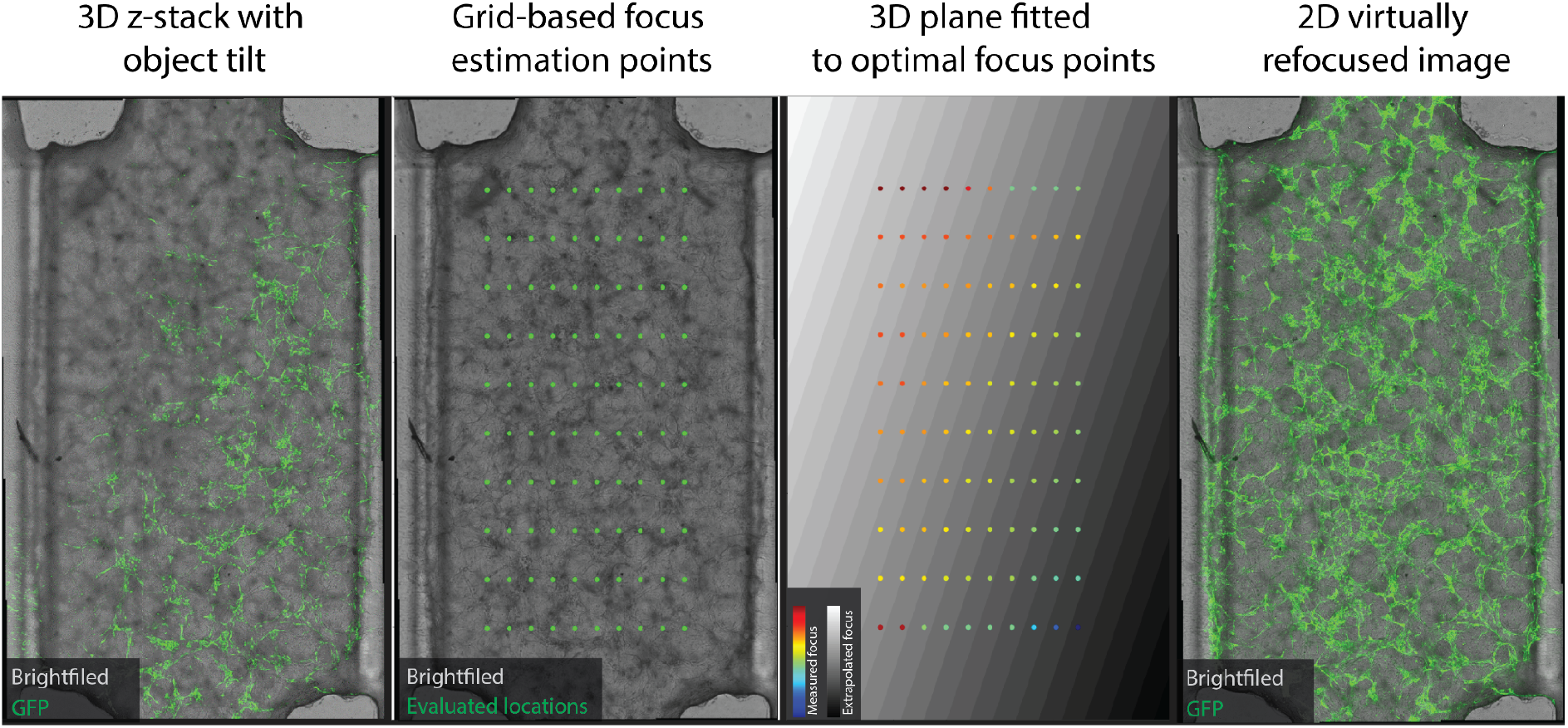
Strategy for virtually refocusing tilted 3D acquisitions. A 3D brightfield microscopy dataset covering the entire vascularized microtissue within the chip was obtained. Due to the planar nature of microvessels and the misalignment with the microscope stage focal plane, microvessels, illustrated by the GFP signal in green, often appear at different heights within each plane, obstructing their complete observation in a single 2D microscope acquisition (first panel). Due to the complex and heterogeneous optical properties of the ECM, existing Z projections adapted to brightfield involving pixel intensity variations are not suitable^20^, consequently a custom virtual refocusing technique was developed to measure the tilted vessel plane within the z-stack. This involved setting focus estimation points (green dots) across the image volume horizontal plane in a grid pattern (second panel). At every point of this grid, patches of 200×200 pixels were defined for each Z plane. The focus of each patch was assessed using the tenengrad variance method, a technique that calculates the variance in the amplitude of the image gradient. The Z plane with the highest tenengrad variance score is deemed the most in-focus for that specific area. After evaluating all potential focal planes across the grid, a three-dimensional plane is aligned to these scores to represent the overall angled focal plane of the vessels (third panel). Finally the initial Z stack is virtually refocused by only sampling its pixels along this tilted plane, resulting in a 2D image that consistently displays the vessels in focus throughout the entire sample (fourth panel). The virtual refocusing was performed using the brightfield channel, the GFP signal in this figure has been included for vessel visualization purposes and was not used for any computations.

**Supplementary Figure 2:**
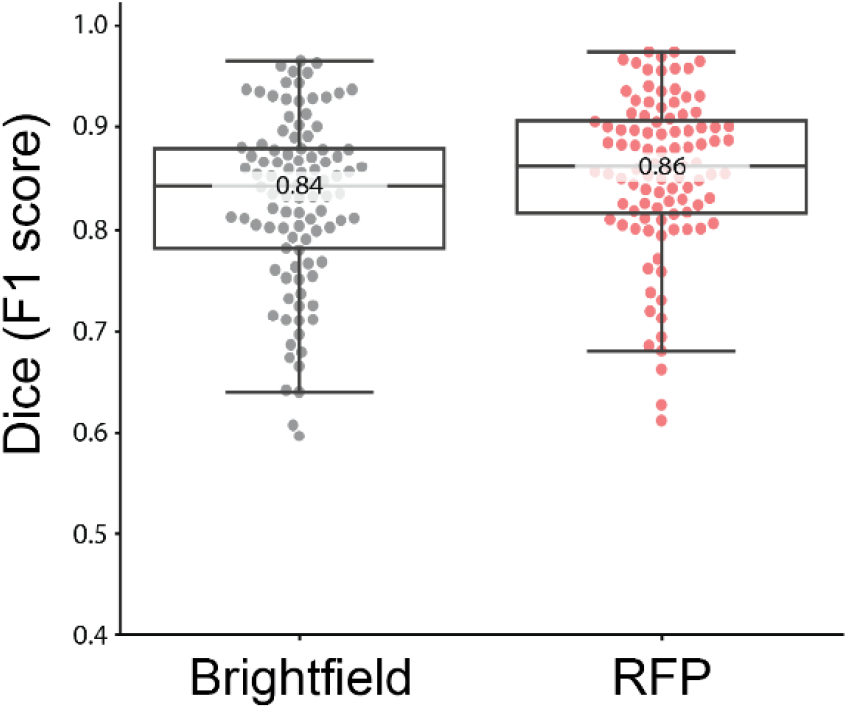
Accurate vessel segmentation in the brightfield modality. A slight improvement in Dice accuracy for vessel segmentation is observed when using fluorescence (red) over brightfield (gray), demonstrating the brightfield as a viable modality for the characterization of vascular networks (two-tailed independent *t*-test p= 0.018, *). Tukey box plots with center line: median, annotated on the figure; box limits, upper and lower quartiles; whiskers, 1.5 × interquartile range.

**Supplementary Figure 3:**
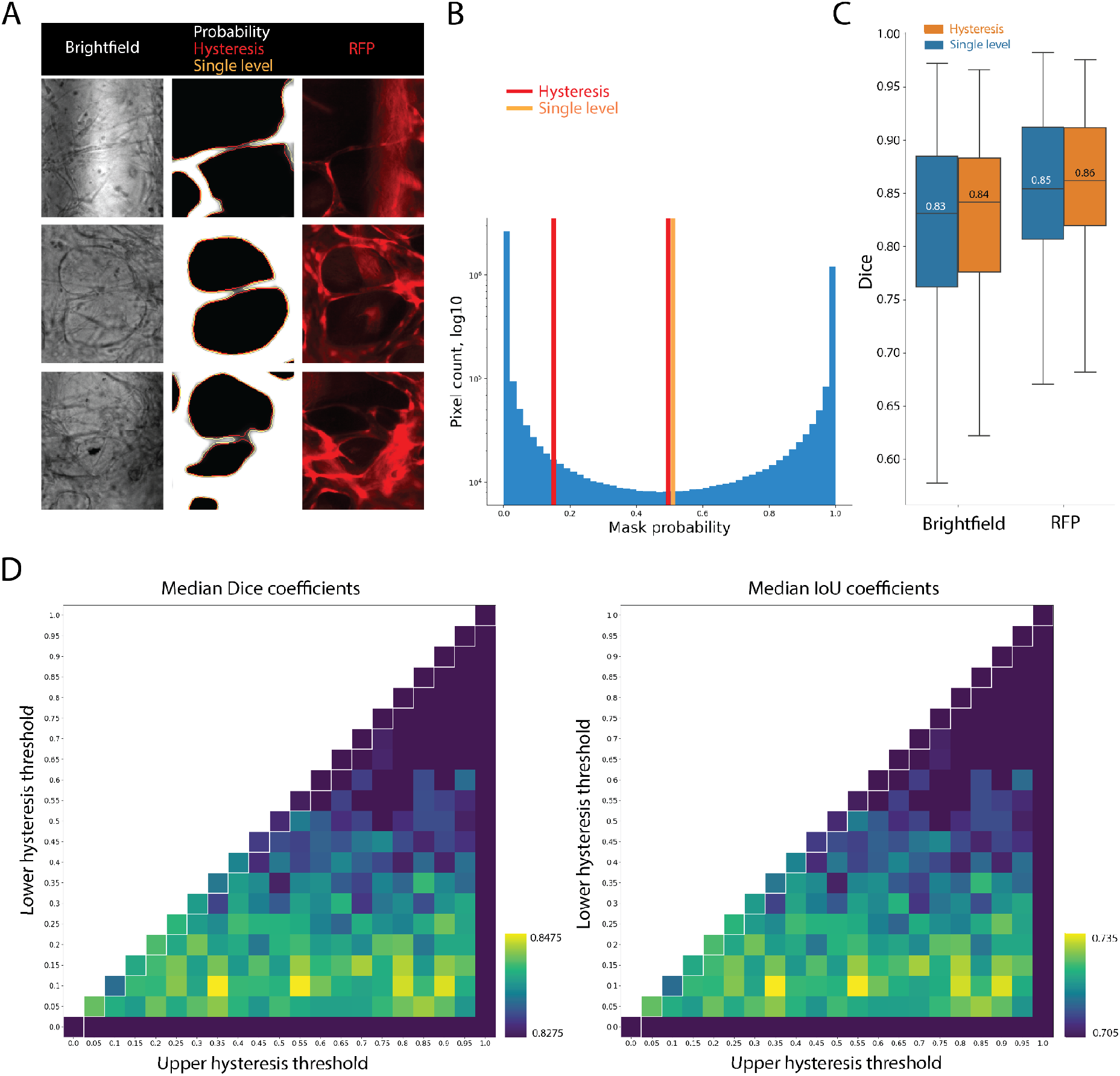
Hysteresis thresholding enables the detection of thin vessels. **A**. Applying hysteresis thresholding (red outline, second column) on the vessel probability map (grayscale, second column) enables the successful classification of thin vessels as foreground, which would otherwise be disconnected if a single-level threshold were applied (orange outline, second column). Hysteresis thresholding employs two thresholds: a lower and an upper limit. Pixels falling below the lower threshold are classified as background, those above the upper threshold as foreground. Pixels with values in between are considered foreground only if directly connected to a pixel exceeding the upper threshold; otherwise, they are classified as background. **B**. The lower and upper hysteresis thresholds have been empirically set at 0.15 and 0.5, respectively (red lines), as explained in D. In contrast, the single-level threshold (orange line) is positioned at the midpoint, 0.5, within the vessel probability map pixel distribution (blue) that spans from 0 to 1. **C**. Dice coefficient distribution of the test set from masks obtained using either hysteresis or single level thresholding, for both the brightfield and fluorescence (RFP) modalities. **D**. Parameter sweep for the lower and upper hysteresis thresholds, reporting either the median Dice or IoU coefficients of the image test set for each threshold combination. Single level thresholding of the vessel probability map was employed (white outlines), at equal values of lower and upper thresholds. While optimal values for the lower threshold seem to fall in the range of 0.05-0.2, the value of the upper threshold does not seem to have a defined optimal range. Therefore, we chose the values of 0.15 and 0.5 for the lower and upper thresholds, respectively.

## Notes

### Competing Interest Statement

The authors have declared no competing interest.

## References

1. Brandt, M. M., Cheng, C., Merkus, D., Duncker, D. J. & Sorop, O. Mechanobiology of Microvascular Function and Structure in Health and Disease: Focus on the Coronary Circulation. Front. Physiol. 12, 771960 (2021).

2. Rizzoni, D. et al. Immune System and Microvascular Remodeling in Humans. Hypertension 79, 691–705 (2022).

3. Beare, J. E., Curtis-Whitchurch, L., LeBlanc, A. J. & Hoying, J. B. Microvasculature in Health and Disease. in Encyclopedia of Cardiovascular Research and Medicine (eds. Vasan, R. S. & Sawyer, D. B.) 349–364 (Elsevier, 2018).

4. Haase, K. & Kamm, R. D. Advances in on-chip vascularization. Regen. Med. 12, 285–302 (2017).

5. Shin, Y. et al. Microfluidic assay for simultaneous culture of multiple cell types on surfaces or within hydrogels. Nat. Protoc. 7, 1247–1259 (2012).

6. Chen, M. B. et al. On-chip human microvasculature assay for visualization and quantification of tumor cell extravasation dynamics. Nat. Protoc. 12, 865–880 (2017).

7. Sontheimer-Phelps, A., Hassell, B. A. & Ingber, D. E. Modelling cancer in microfluidic human organs-on-chips. Nat. Rev. Cancer 19, 65–81 (2019).

8. Tomanek, R. J. The coronary capillary bed and its role in blood flow and oxygen delivery: A review. Anat. Rec. 305, 3199–3211 (2022).

9. Salipante, P. F., Hudson, S. D. & Alimperti, S. Blood vessel-on-a-chip examines the biomechanics of microvasculature. Soft Matter 18, 117–125 (2021).

10. Masciantonio, M. G., Lee, C. K. S., Arpino, V., Mehta, S. & Gill, S. E. Chapter Three - The Balance Between Metalloproteinases and TIMPs: Critical Regulator of Microvascular Endothelial Cell Function in Health and Disease. in Progress in Molecular Biology and Translational Science (ed. Khalil, R. A.) vol. 147 101–131 (Academic Press, 2017).

11. Zudaire, E., Gambardella, L., Kurcz, C. & Vermeren, S. A computational tool for quantitative analysis of vascular networks. PLoS One 6, e27385 (2011).

12. Kirst, C. et al. Mapping the Fine-Scale Organization and Plasticity of the Brain Vasculature. Cell 180, 780–795.e25 (2020).

13. Spangenberg, P. et al. Rapid and fully automated blood vasculature analysis in 3D light-sheet image volumes of different organs. Cell Rep Methods 3, 100436 (2023).

14. Vickerman, M. B. et al. VESGEN 2D: automated, user-interactive software for quantification and mapping of angiogenic and lymphangiogenic trees and networks. Anat. Rec. 292, 320–332 (2009).

15. Seaman, M. E., Peirce, S. M. & Kelly, K. Rapid analysis of vessel elements (RAVE): a tool for studying physiologic, pathologic and tumor angiogenesis. PLoS One 6, e20807 (2011).

16. Corliss, B. A. et al. REAVER: A program for improved analysis of high-resolution vascular network images. Microcirculation 27, e12618 (2020).

17. Akinbote, A. et al. Classical and Non-classical Fibrosis Phenotypes Are Revealed by Lung and Cardiac Like Microvascular Tissues On-Chip. Front. Physiol. 12, 735915 (2021).

18. Moccia, C. et al. Mammary Microvessels are Sensitive to Menstrual Cycle Sex Hormones. Adv. Sci. e2302561 (2023).

19. Rota, A. et al. A three-dimensional method for morphological analysis and flow velocity estimation in microvasculature on-a-chip. Bioeng. Transl. Med. (2023) doi:10.1002/btm2.10557.

20. Selinummi, J. et al. Bright field microscopy as an alternative to whole cell fluorescence in automated analysis of macrophage images. PLoS One 4, e7497 (2009).

21. Implementation of Automatic Focusing Algorithms for a Computer Vision System with Camera Control. The Robotics Institute Carnegie Mellon University https://www.ri.cmu.edu/publications/implementation-of-automatic-focusing-algorithms-for-a-computer-vision-system-with-camera-control/ (1983).

22. Ronneberger, O., Fischer, P. & Brox, T. U-Net: Convolutional Networks for Biomedical Image Segmentation. arXiv [cs.CV] (2015).

23. Cherubini, M. et al. Flow in fetoplacental-like microvessels in vitro enhances perfusion, barrier function, and matrix stability. Sci Adv 9, eadj8540 (2023).

24. Douglas, S. A., Haase, K., Kamm, R. D. & Platt, M. O. Cysteine cathepsins are altered by flow within an engineered in vitro microvascular niche. APL Bioeng 4, 046102 (2020).

25. Haase, K., Gillrie, M. R., Hajal, C. & Kamm, R. D. Pericytes Contribute to Dysfunction in a Human 3D Model of Placental Microvasculature through VEGF-Ang-Tie2 Signaling. Adv. Sci. 6, 1900878 (2019).

26. Weigert, M. et al. Content-aware image restoration: pushing the limits of fluorescence microscopy. Nat. Methods 15, 1090–1097 (2018).

27. Aguet, F., Van De Ville, D. & Unser, M. Model-based 2.5-d deconvolution for extended depth of field in brightfield microscopy. IEEE Trans. Image Process. 17, 1144–1153 (2008).

28. Wu, Y. et al. Three-dimensional virtual refocusing of fluorescence microscopy images using deep learning. Nat. Methods 16, 1323–1331 (2019).

